## Supplemental Materials for "A hepatitis B virus (HBV) sequence variation graph improves sequence alignment and sample-specific consensus sequence construction for genetic analysis of HBV"

**Supplementary Materials**

**HBV reference graph** (doi: 10.5281/zenodo.6646207)

**Supplementary Figures**

**Figure S1**: Sequence-to-graph alignment

**Figure S2:** Distribution of path depth across nodes within the HBV reference graph.

**Figure S3**: Alignment comparisons of simulated HBV genotype B/C sequencing data.

**Figure S4:** Proportion of successfully aligned CHB sequencing data.

**Figure S5**: Distribution of rescued linear alignments.

**Figure S6**: Distribution of rescued graph alignments.

**Figure S7:** Comparison of linear reference-based alignment approaches.

**Figure S8**: Mash distance comparisons of consensus sequences: simulated HBV sequencing data.

**Figure S9**: Mash distance comparisons of consensus sequences: simulated HBV genotypes B/C sequencing data.

**Figure S10**: Genetic distance comparisons of consensus sequences and de novo assembled HBV haplotypes with CHB sequencing data from longitudinal CHB samples.

**Supplementary Tables**

**Table S1**: Computational time comparisons of graph and linear reference-based alignment.

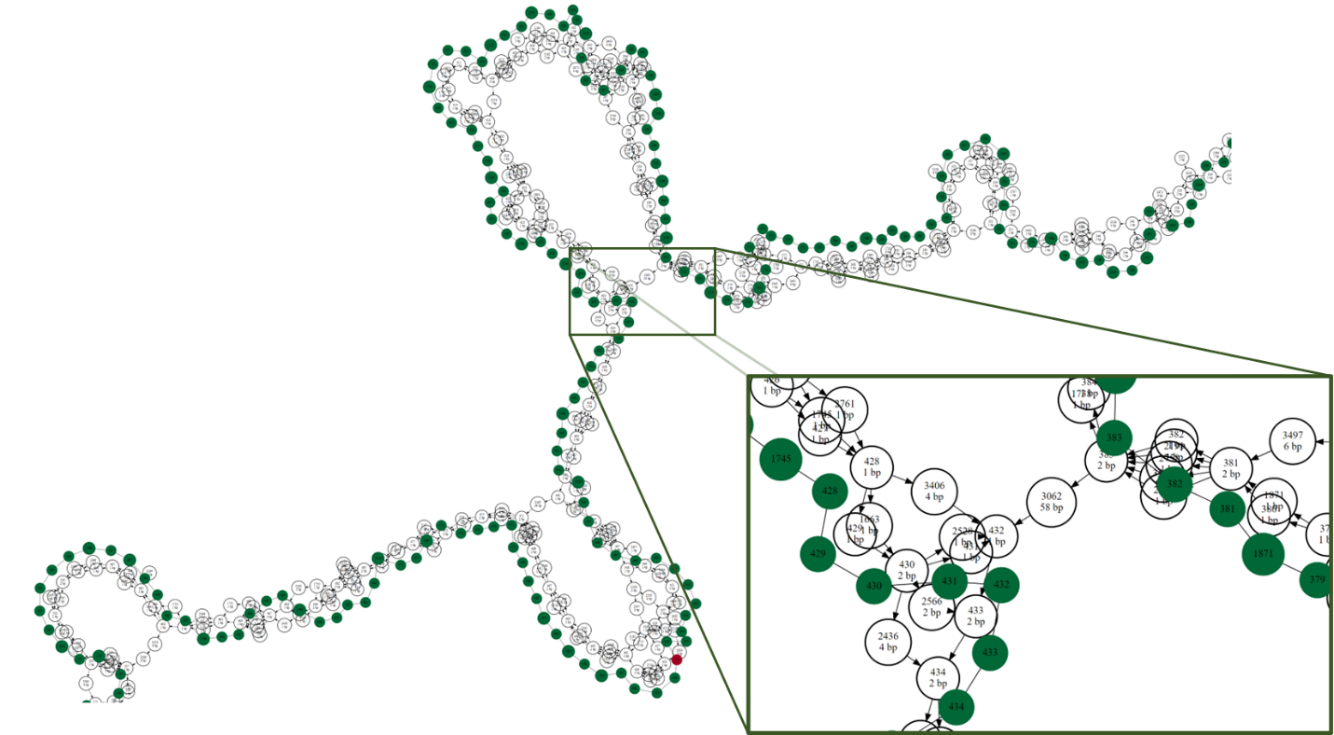

**Figure S1**: Sequence-to-graph alignment. Alignment of a full-length HBV genome sequence (GenBank ID: X52939,^127^ HBV genotype C) to a localized region of a genome graph constructed from the 44 HBV genomes used to construct the reference graph.^120^ The square panel reflects an enlarged subregion of the graph alignment to aid visualization of the sequence-to-graph alignment. Nodes reflect stretches of nucleotide sequence which are connected to one another via edges, with the sequence length of each node provided beneath unique node identifiers. The HBV reference graph is comprised of the colorless nodes, while nodes to which the sequence successfully aligns are repeated and colored from green to red, reflecting good to bad sequence-to-node matching quality, respectively.

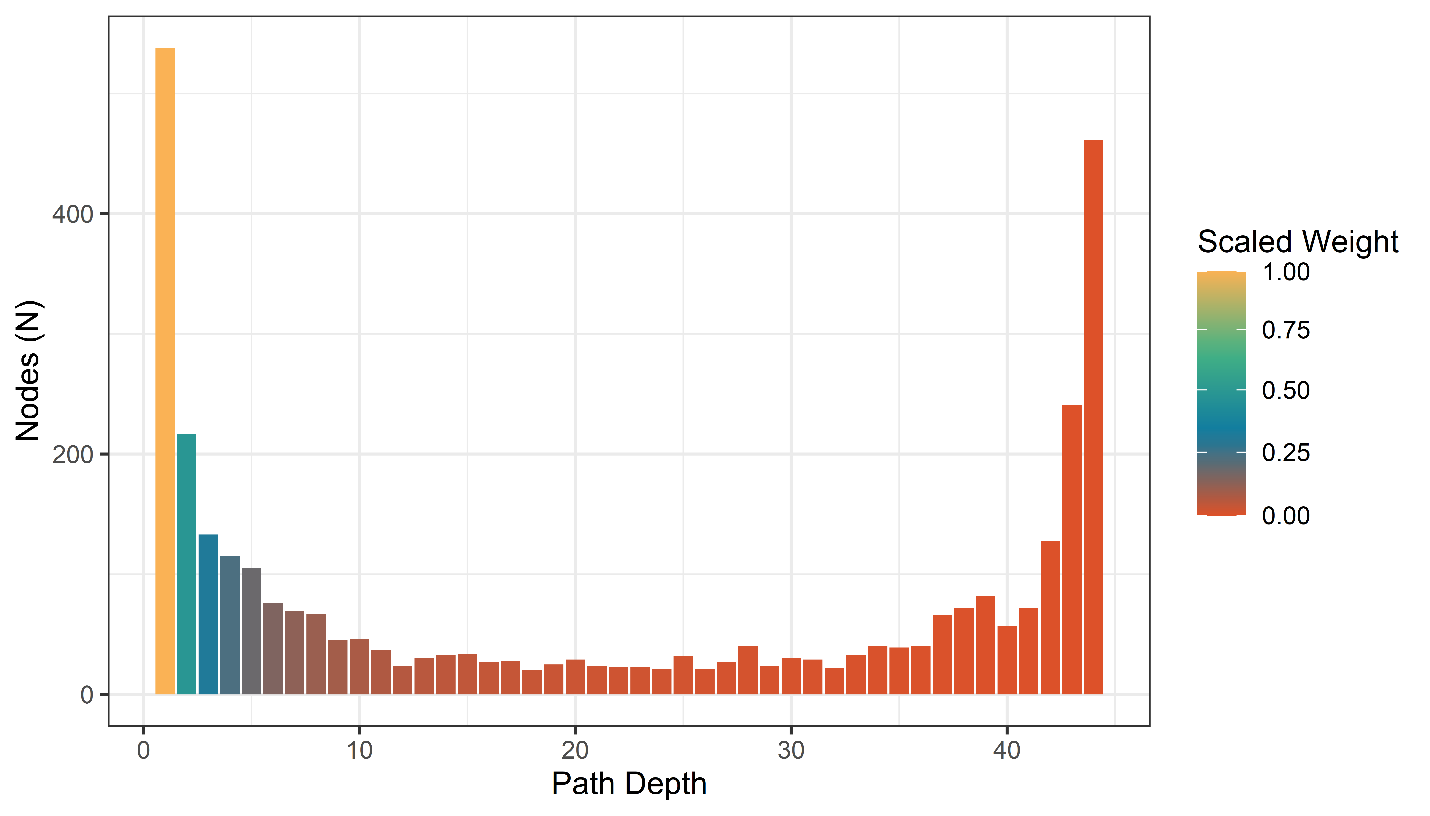

**Figure S2**: Distribution of path depth across nodes within the HBV reference graph. The Y axis and heigh of each bar reflects the number of nodes across the HBV reference graph with a specific path depth. The X axis reflects path depth, or the number of paths traversing a single node. Colors of the bar indicate the scaled weight associated with path depth, with nodes unique to a single path/HBV genome sequence having the maximum weight and nodes traversed by every path/genome weighted to 0.

**Figure S3**: Alignment comparisons of simulated HBV genotype B/C sequencing data. Points reflect the proportion of successfully aligned sequences, colored by either the genotype of the reference used for linear reference-based alignment or whether graph-based alignment was performed. The Y axis reflects the proportion of successfully aligned reads. Labels reflect the genotype of the reference sequence or graph used in the alignment and the proportion of reads successfully aligned. The right panel reflects alignment of HBV sequencing data generated from HBV genotype B or C sequences, highlighting the highest and lowest proportion of reads aligned to references of each genotype, along with the proportion of reads successfully aligned to the HBV reference graph.

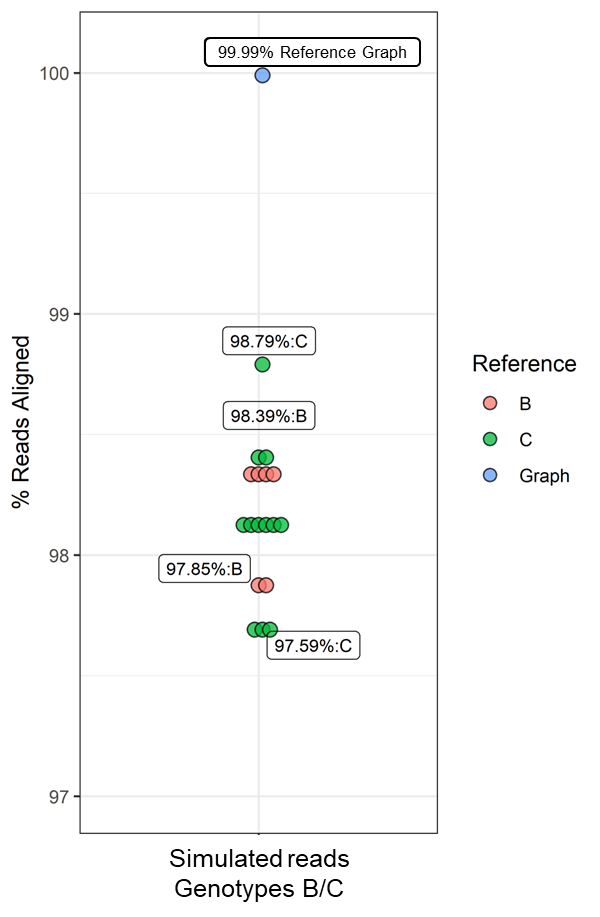

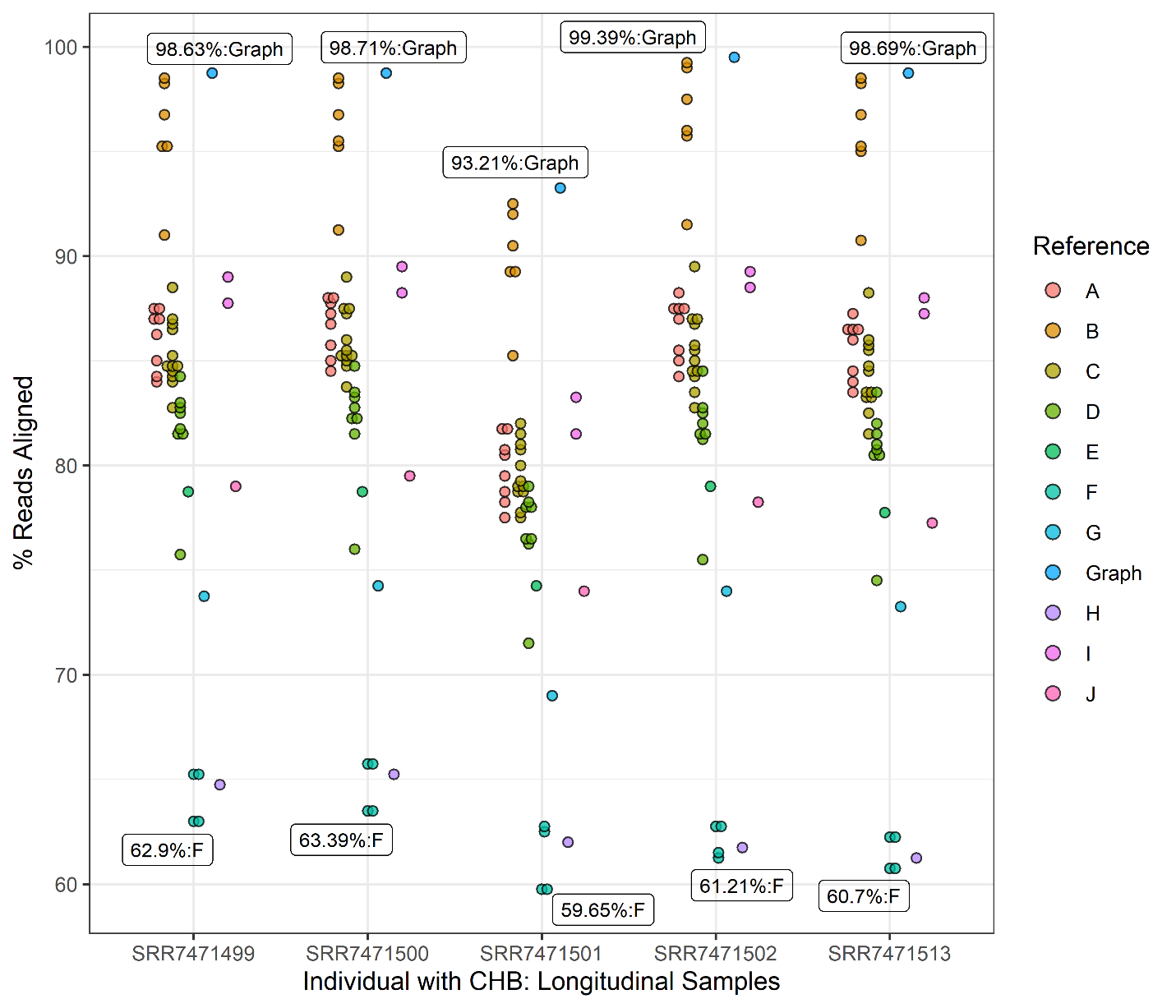

**Figure S4**: Proportion of successfully aligned CHB sequencing data. Points reflect the proportion of successfully aligned sequences, colored by either the genotype for linear reference-based alignment or if sequences were aligned to the HBV reference graph. The Y axis reflects the proportion of successfully aligned reads. The X axis indicates each longitudinally collected CHB sample. Labels at the highest and lowest observed aligned proportions reflect the reference’s genotype or whether a graph-based alignment was performed.

**Figure S5**: Distribution of rescued linear alignments. Sub-optimal linear-based alignments each had their unmapped reads re-aligned to the best performing linear reference (HBV subgenotype B2). The X axis reflects HBV genome position. The Y axis reflects a scaled proportion of successful re-alignments. Colored and labeled lines indicate the genotype of the initial reference used for sequence alignment. Gene-encoding regions have been provided below each figure. C=core, P=polymerase, S=Surface, X=X.

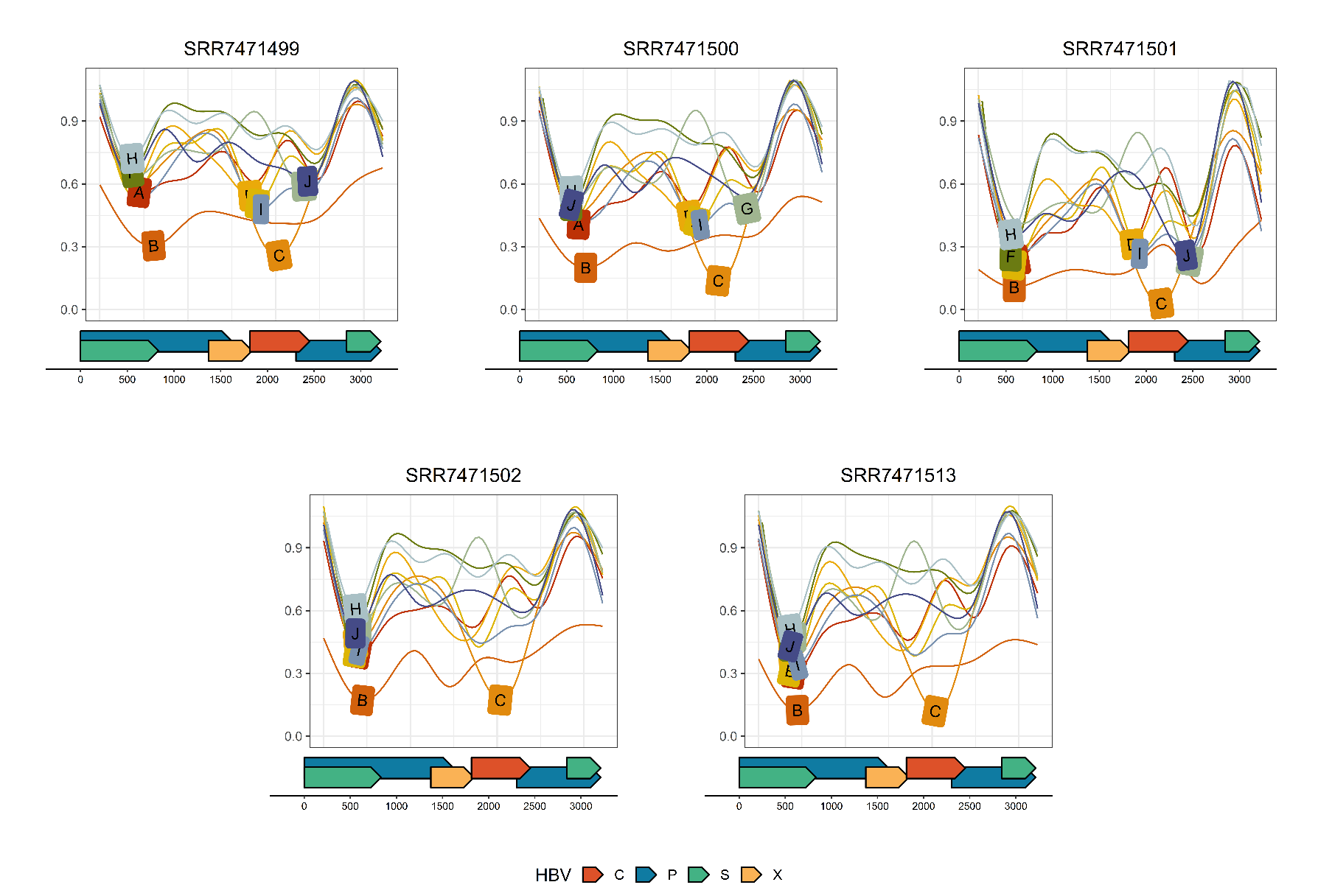

**Figure S6**: Distribution of rescued graph alignments. Sequences unable to align to linear reference sequences of the best performing HBV subgenotype (B2) were aligned to the HBV reference graph. The very top panel reflects a Bandage visualization of the compacted de Bruijn Graph created using the 44 proposed HBV reference sequences. The middle panel reflects the average nucleotide diversity across the genome, with the Y axis reflecting the average pairwise nucleotide diversity (0-1). For all panels, the X axis reflects genome position and for the bottom panel, the Y axis reflects the density of the start sites observed for each successful sequence-to-graph alignment. Colored areas beneath density curves reflect sample-specific alignment density (legend embedded within bottom panel). Gene regions are provided at the base of the figure. C=core, P=polymerase, S=Surface, X=X.

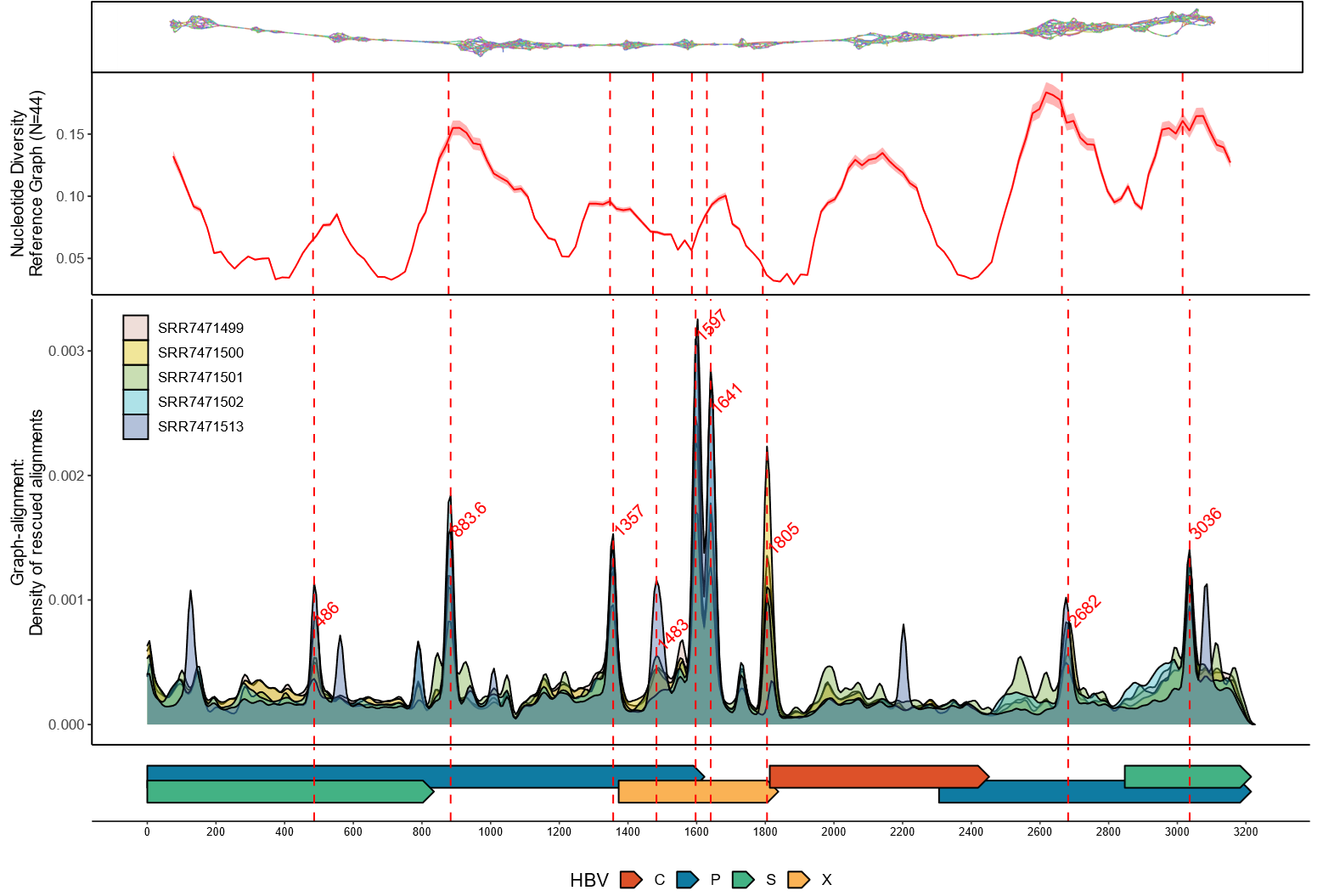

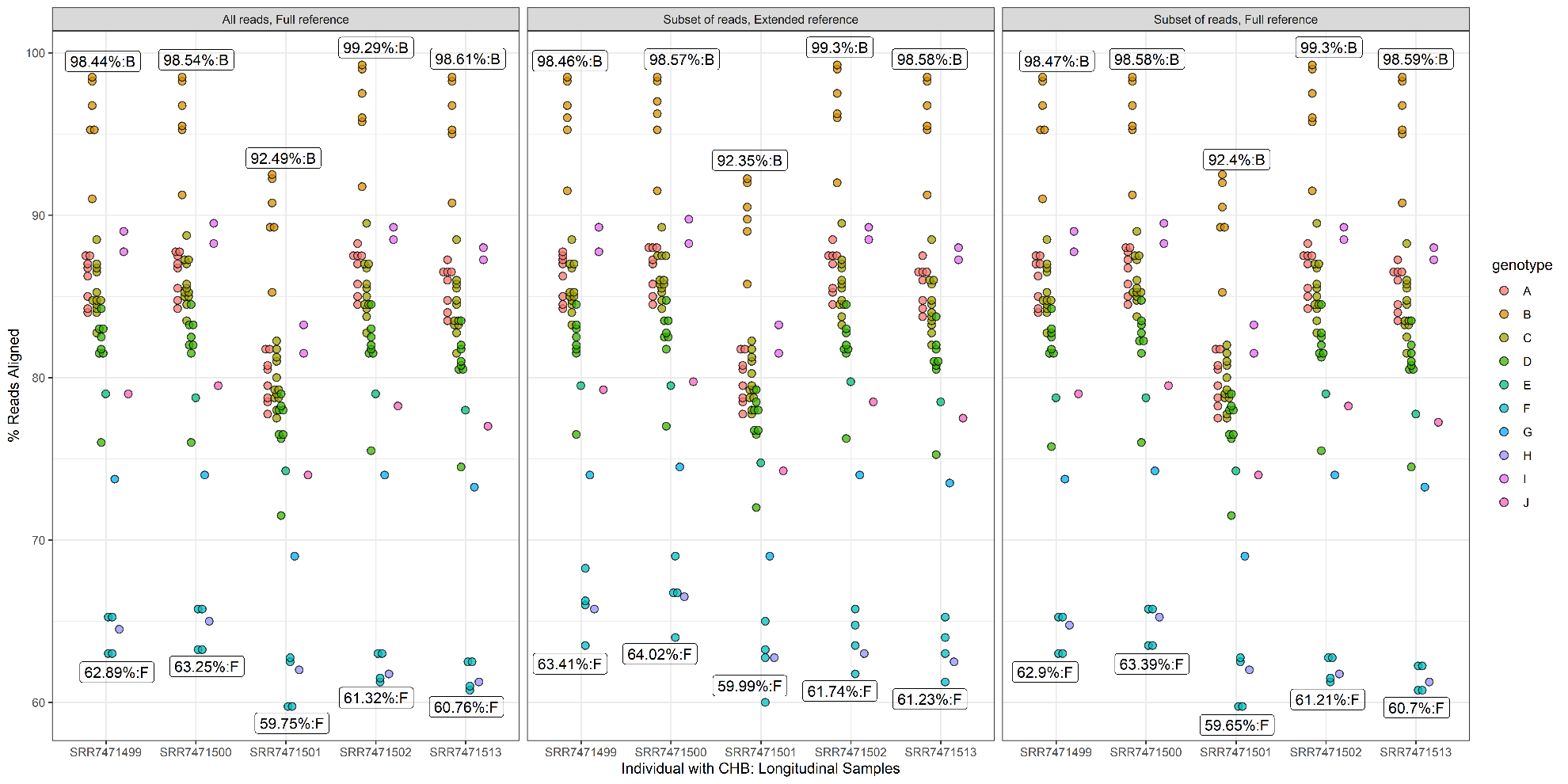

**Figure S7:** Comparison of linear reference-based alignment approaches. Points reflect the proportion of successfully aligned sequences observed when all QC-passed CHB sequencing data was used with the full-length reference sequence (left), when a subsampled dataset was used with an extended reference sequence (middle), and when a subsampled dataset was used with the full-length reference sequence (right). Points are colored by the genotype of the linear reference sequence used in alignment. The Y axis reflects the proportion of successfully aligned reads. The X axis indicates CHB sample. Labels at the highest and lowest observed proportions also indicate the genotype of the reference used in alignment.

**Figure S8**: Mash distance comparisons of consensus sequences: simulated HBV sequencing data. Consensus sequences generated from alignments to 44 phylogenetically representative linear HBV reference sequences or the HBV reference graph. The Y axis reflects the Mash distance estimated between each consensus sequence and the set of 59 full-length HBV genome sequences used to simulate the reads.

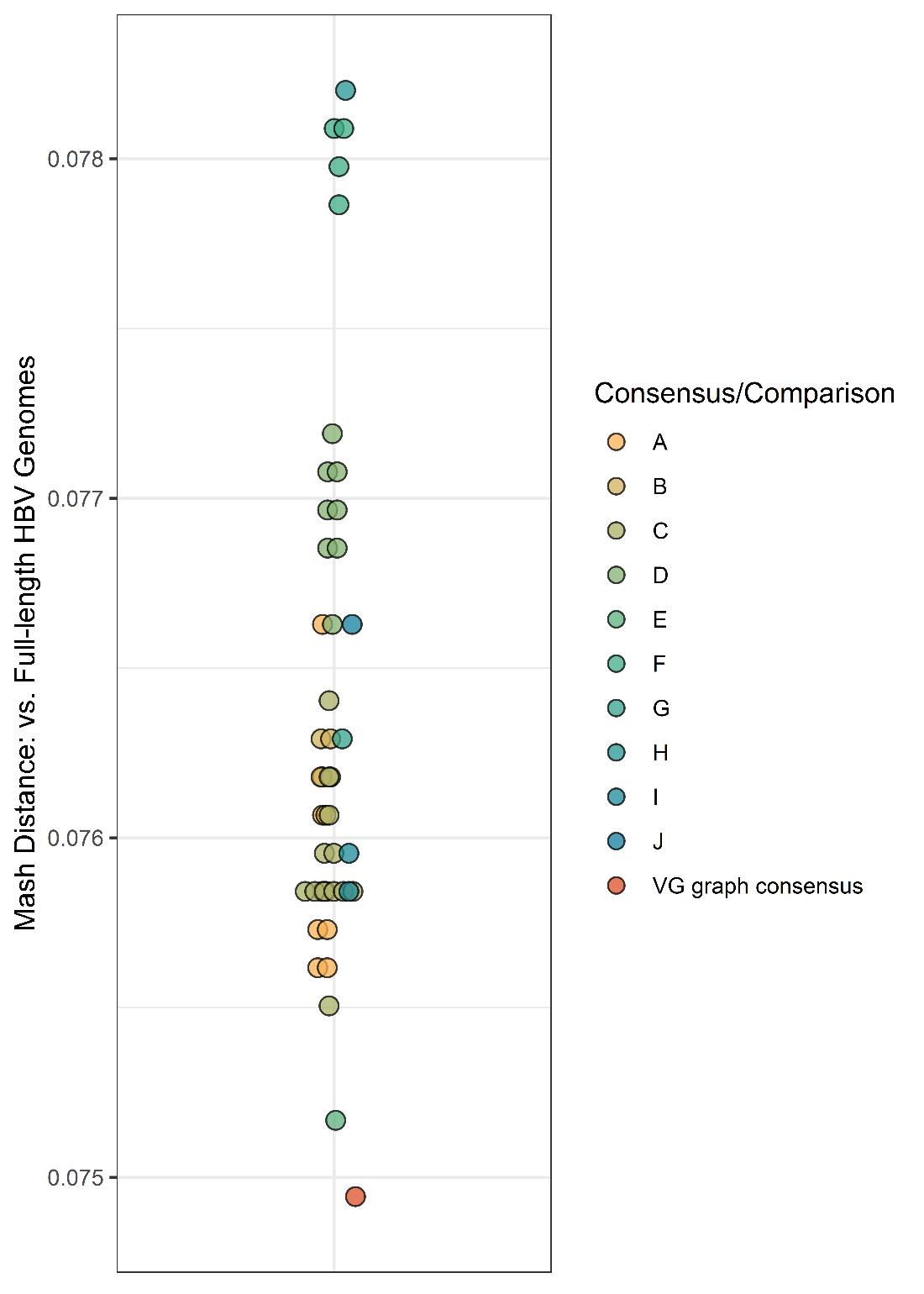

**Figure S9**: Mash distance comparisons of consensus sequences: simulated HBV genotypes B/C sequencing data. Consensus sequences generated from HBV genotype B/C sequencing data aligned to reference sequences from genotypes B/C or the HBV reference graph. The Y axis reflects the Mash distance estimated between each consensus sequence and the full-length HBV genotype B/C sequences used to simulate the reads (N=18).

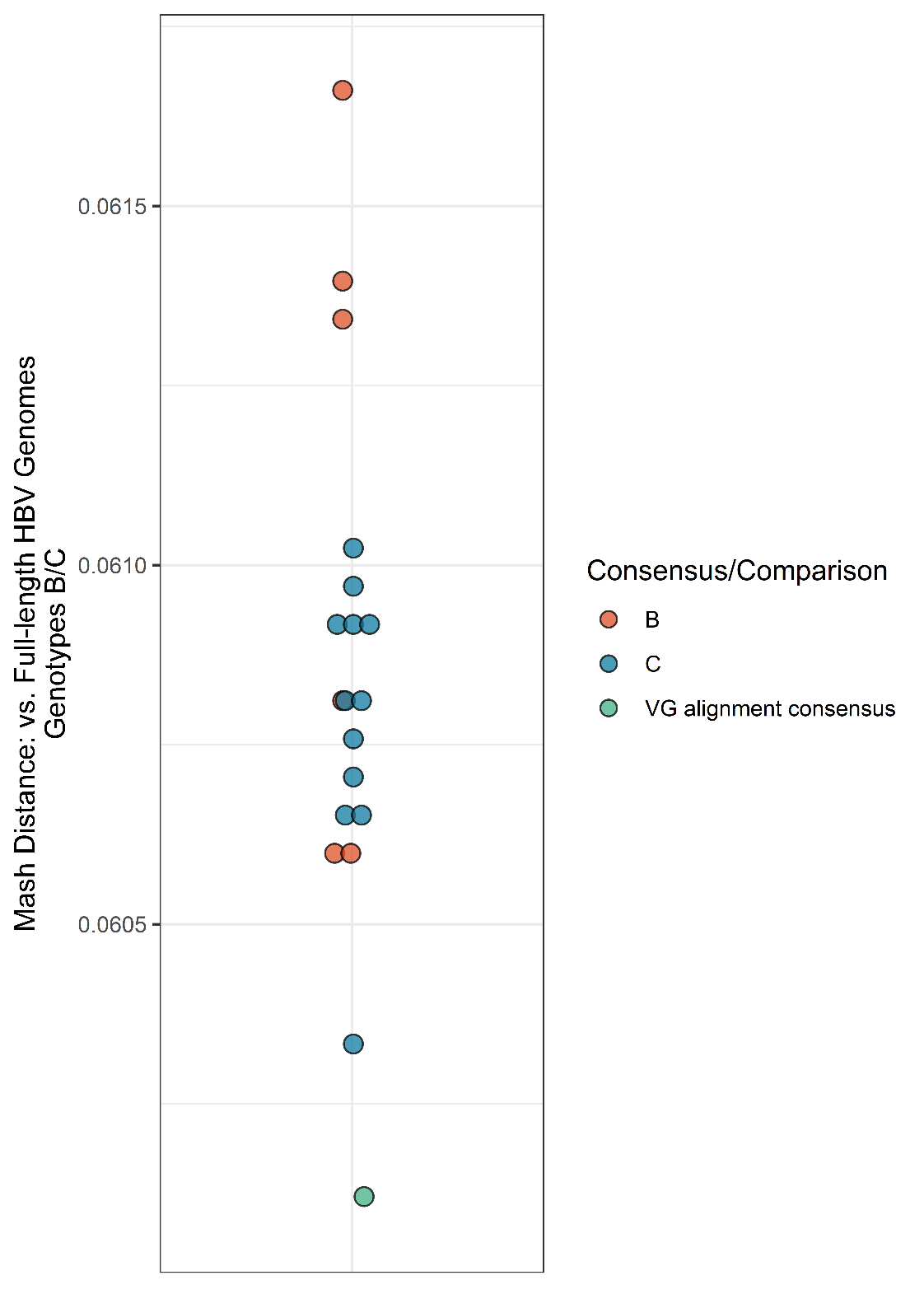

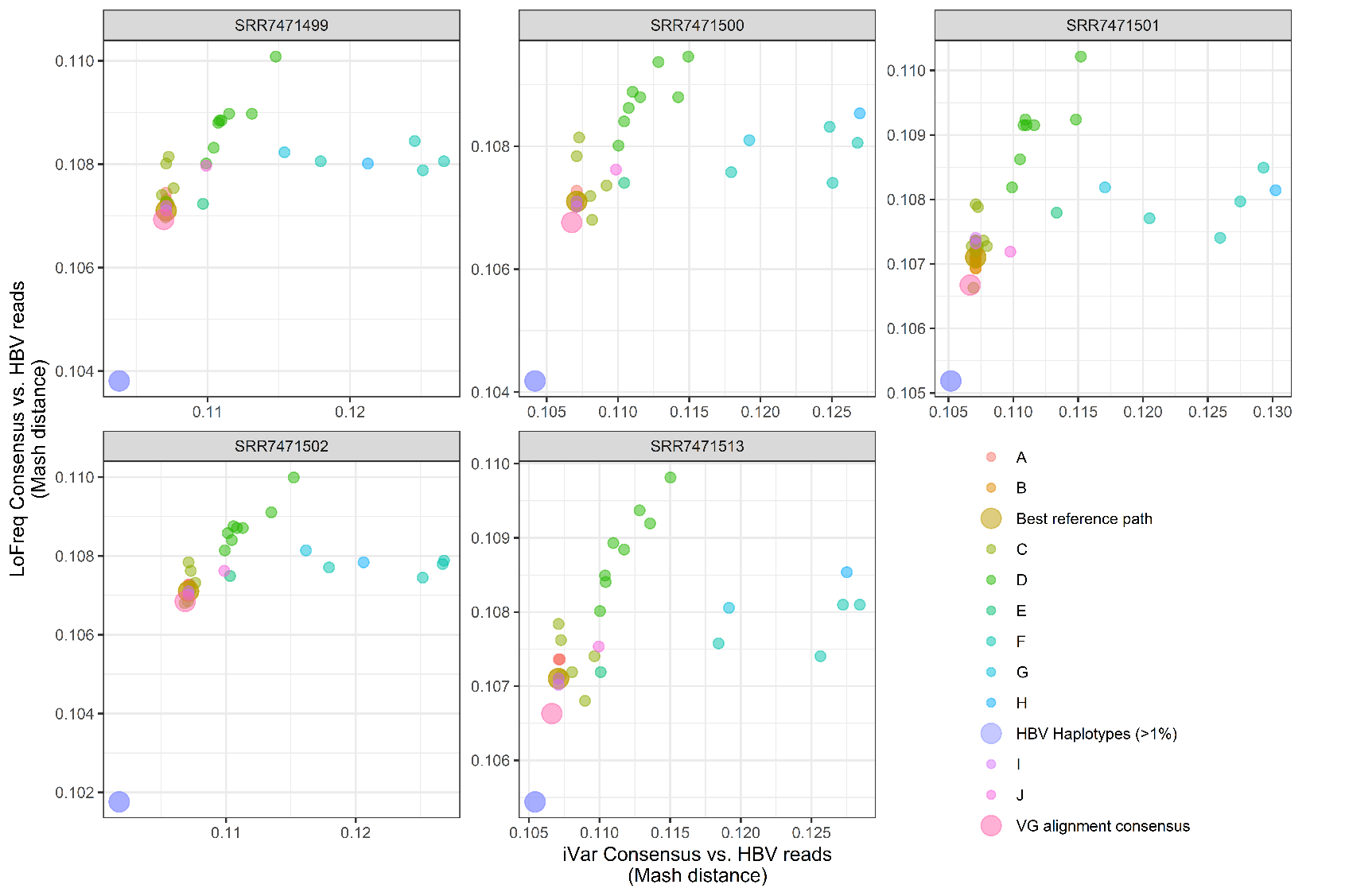

**Figure S10**: Genetic distance comparisons of consensus sequences and de novo assembled HBV haplotypes with CHB sequencing data from longitudinal CHB samples. Points reflect the Mash distance estimated between each consensus sequence generated from the 44 HBV reference sequences or the HBV reference graph for each of the five longitudinally collected CHB samples. The Y axis reflects the Mash distance estimated between each LoFreq-derived consensus sequence and the X axis reflects the Mash distance estimated between each iVar-derived consensus sequence. The color of each point reflects the genotype of the reference used to generate a consensus, or if the consensus was derived via graph-based alignment or reflects sample-specific HBV haplotypes. Points for graph-derived consensus sequences, including VG-based variant calling (‘VG alignment consensus’), graph-based surjection (‘Best reference path’), and the de novo assembled viral strains (‘HBV Haplotypes (>1%)’) are enlarged.

| **Time metric** | **BWA-MEM** | **VG Giraffe** | **VG Giraffe (fast)** | **VG Map** |
| --- | --- | --- | --- | --- |
| Wall clock time | 25.0 | 11.6 | 6.6 | 877.6 |
| CPU time (total) | 53.2 | 57.0 | 56.6 | 101,432.9 |
| CPU time (user-mode) | 50.9 | 53.9 | 54.0 | 101,367.3 |
| CPU time (system) | 2.3 | 3.0 | 2.6 | 65.6 |

**Table S1**: Computational time comparisons of graph and linear reference-based alignment. Wall clock time and CPU time were measured using /usr/bin/time and are presented in minutes. Time reflects the resources required to align simulated high diversity HBV sequences to either 44 linear reference sequences or a graph comprised of these sequences followed by the estimation of alignment statistics using either SAMtools or VG, respectively. All mapping was performed using 16 threads.
